# Plasticity in Ligand Recognition at Somatostatin Receptors

**DOI:** 10.1101/2021.11.02.466988

**Authors:** Michael J. Robertson, Justin G. Meyerowitz, Ouliana Panova, Kenneth Borrelli, Georgios Skiniotis

## Abstract

Somatostatin is a signaling peptide that plays a pivotal and wide-ranging role in physiologic processes relating to metabolism and growth through its actions at somatostatin receptors (SSTRs). Members of the somatostatin receptor subfamily, particularly SSTR2, are key drug targets for neuroendocrine neoplasms, with several synthetic peptide agonists currently in clinical use. Here, we show the cryogenic electron microscopy (cryo-EM) structures of active-state SSTR2 in complex with heterotrimeric Gi protein and either the endogenous ligand SST14 or the FDA-approved drug octreotide.

Complemented by biochemical assays and molecular dynamics simulations, these structures reveal key details of ligand recognition, receptor activation, and subtype-selectivity at somatostatin receptors. We find that SSTR ligand recognition is highly diverse, as demonstrated by ligand-induced conformational changes in ECL2, substantial sequence divergence across subtypes in extracellular regions, and loss of ligand binding upon several structurally homologous substitutions between subtypes. Despite this complexity, we rationalize several previously described sources of SSTR subtype selectivity and identify an additional key interaction for SSTR2/3/5 specific binding. These results shed light on the basis of ligand recognition by somatostatin receptors and provide valuable insights for structure-based drug discovery at these targets.

## Introduction

Somatostatin (SST) is a 14- (SST14) or 28-residue (SST28) peptide that regulates crucial aspects of animal physiology by suppressing metabolic- and growth-related neuroendocrine signaling systems including insulin/glucagon, thyroid-stimulating hormone, prolactin, growth hormone, and digestion-related processes^1^. These actions of somatostatin are mediated by somatostatin receptors (SSTRs), which are G protein-coupled receptors (GPCRs) that signal through the inhibitory Gi/o family of G proteins^2^ to prevent the release of secretory hormone vesicles. Neuroendocrine tumors (NETs) generally overexpress SSTRs, allowing somatostatin agonists to function as diagnostic imaging tracers^3^ and therapeutic agents for hormone excess disorders caused by NETs, including carcinoid syndrome, gigantism, acromegaly, hyperthyroidism, and Cushing’s disease^4^.

The SSTR subfamily includes five distinct isoforms, SSTR1-5, with SSTR2 having two C-terminal splice variants, SSTR2a and SSTR2b. These receptors have varied tissue expression profiles, including differential expression across NETs. SSTR2 is highly overexpressed in most NETs, and thus this subtype is the primary target for clinical agonists. All FDA-approved somatostatin agonists currently in use are peptide analogs of SST14, including the SSTR2-selective drugs octreotide and lanreotide^5^, and the newer SSTR2-, 3-, and 5-selective pasireotide^6^. Abundant evidence suggests that distinct SSTR isoforms play important roles in a variety of tumor subtypes beyond NETs^1^, elevating the need to probe isoform-specific function with more selective pharmacologic tools^7^. Furthermore, the development of orally bioavailable small-molecule SSTR agonists could substantially improve treatment of NETs, as these agonists are often required for chronic therapy^8^. However, the lack of structural information for either the inactive or active state of any of the SSTR subtypes, and the lack of a structural framework for SSTR ligand binding and selectivity, is a substantial impediment for further drug discovery.

All SSTR subtypes exhibit nearly identical affinity for the endogenous peptide SST14, despite having only 40-55% sequence homology and substantial variation in the extracellular region encompassing the ligand interaction site. Radioligand binding assays have suggested that both the extracellular half of the transmembrane (TM) domains and the extracellular loops (ECLs) play a role in ligand binding and subtype selectivity^9^. However, which precise residue(s) are critical depends upon both subtype and ligand, suggesting a complex interaction landscape between SSTRs and their ligands. Here, we used cryo-EM, biochemical assays, and atomistic simulations to obtain structural and mechanistic insights into ligand recognition and selectivity at somatostatin receptors. The results highlight surprising heterogeneity and diversity of ligand-protein recognition in a subfamily of receptors that are activated by the same endogenous ligand.

### SSTR2 Couples Robustly to Gi/o Proteins

To better understand the signaling landscape of SSTR2 and identify the optimal G protein partner for forming stable receptor-G protein complexes for structural studies we employed bioluminescence resonance energy transfer (BRET)-based activation assays to screen for coupling by the majority of G proteins^10^ (Extended Data Fig. 1a, b). Consistent with previous studies, SSTR2 interacted robustly with all Gi/o family G proteins. Although Gi1 has traditionally been the preferred Gi/o type for structural studies, our results identified it as the weakest SSTR2 signaling partner within the Gi/o family. Interestingly, we also observed ligand-induced interaction with G15, which belongs to the Gq family, and G12, which forms a distinct G protein subfamily with G13, although in both cases with weak potency compared to Gi/o activation. No other G proteins tested of the Gq, Gs, or G12/13 families were activated by SSTR2. As prior studies have established Gi3 as a likely physiologic coupling partner, we used this G protein for complexation and further studies. To resolve receptor stability issues at low ionic strength during purification, we swapped the third intracellular loop (ICL3) and the bottom 9 residues of transmembrane helix 6 (TM6) for that of the kappa opioid receptor (KOR)^11^. This strategy also enabled us to employ a nanobody recognizing the KOR ICL3 in order to determine the inactive-state cryo-EM structure of SSTR2 reported in a companion manuscript (Robertson et al.^12^). This construct, denoted SSTR2_κICL3_, coupled to Gαi3 indistinguishably from wild-type receptor in BRET assays (Extended Data Fig. 1c). Thus, we used SSTR2_κICL3_ for complexation with a dominant-negative Gαi3βγ heterotrimer and scFv16, an antibody fragment that aids complex stability and has been employed in numerous cryo-EM studies^13,14^. High-quality complex preparations were obtained with both octreotide and SST14, and cryo-EM enabled us to determine maps with global resolutions of 2.9 Å and 2.5 Å, respectively (Fig. 1a, Extended Data Fig. 2, 3). The two structures show near-identical coupling between the receptor and Gαi3, in a way that is similar to the receptor-G protein coupling interaction of the μ-opioid receptor in complex with Gαi1^15^ (Extended Data Fig. 4a). At the resolutions obtained, water-mediated interactions can be resolved between the α5 helix of Gαi3 and the intracellular cavity of the receptor, bridging the TM3 DRY motif D139^3.49^ (Ballesteros-Weinstein notation^16^), R140^3.50^, T78^2.39^, R155^4.38^, S150 and the backbone carbonyl of P147 on the receptor with E350 N347 and the backbone carbonyl of C351 on the G protein (Extended Data Fig. 4b, c). These density interpretations were further confirmed by JAWS simulations^17^, which probe occupancies and absolute binding free energies of water molecules, that recapitulated the experimentally observed water positions at the receptor-G protein interface (Extended Data Fig. 4b, c).

**Figure 1.**
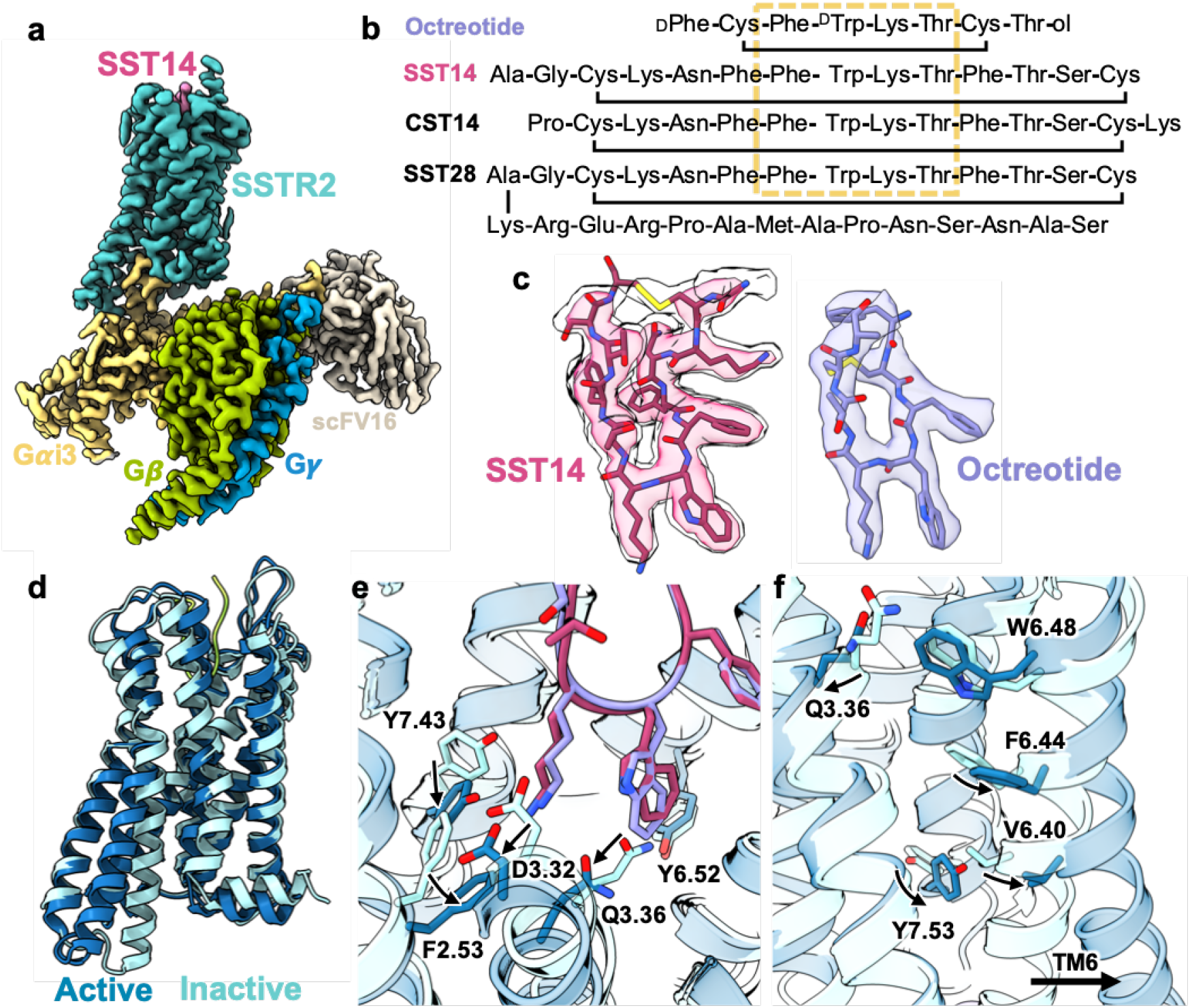
Cryo-EM structures of SST14 or octreotide activated SSTR2-Gi protein complex. **a,** Active-state SSTR2 (teal) bound to SST14 (magenta) in complex with G**α**i3 (goldenrod), G**β** (lime), G**γ** (cerulean), and scFv16 (bone). **b,** Peptide agonists of SSTR2, with conserved key binding region boxed in goldenrod. **c,** map-model fit of SST14 (magenta, left) at two different threshold levels and octreotide (lavender, right). **d,** Alignment of active (blue) and inactive (arctic blue) SSTR2 receptors. **e,** Peptide-induced conformational changes in the extracellular cavity during activation in response to SST14 (magenta) or octreotide (lavender). **f,** Conformational changes on the intracellular site of SSTR2 upon ligand binding and G protein coupling.

### Agonist-Bound SSTR2 Structures Reveal Flexible Ligand Accommodation

Both SST14 and octreotide were well-resolved in the orthosteric binding site, with the conserved Phe-^L/D^Trp-Lys-Thr motif, common to most SSTR-binding peptides, located in the core of the receptor (Fig 1b, c, e). Comparing the ligand-bound structures to our apo, inactive-state structure (Fig. 1d), the SST14/octreotide tryptophan buries into a hydrophobic pocket, breaking a hydrogen bond between Q126^3.36^ and Y273^6.52^ that is present in the apo structure (Fig. 1e). The lysine (L5 of octreotide and L9 of SST14) forms a salt bridge with D122^3.32^, causing this aspartate and Y273^6.52^ to shift position, which in turn drives F92^2.53^ to shift down compared to the inactive state, pushing against TM3. The combination of the steric pressure from F92^2.53^ and the loss of the Q126^3.36^-Y273^6.52^ hydrogen bond allows TM3 to be displaced away from TM6 by roughly 3.5 Å. This movement creates the necessary space for the PIF-motif phenylalanine (F267^6.46^)^18^ to alter position and Y312^7.53^ of the NPxxY motif^19^ to engage in TM6 opening and activation of the receptor (Fig. 1f). The Q126^3.36^-Y273^6.52^ TM3/TM6 hydrogen bond motif is somewhat unique, with only SSTR2, SSTR3, SSTR5, and the melanin-concentrating hormone receptors 1 and 2 (MCH1/2) having a Q^3.36^, and only SSTR2, SSTR3, and MCH1 possessing Q^3.36^ paired with Y^6.52^. Of note, W269^6.48^, often cited as a ‘toggle switch’ residue that changes conformation upon activation of several family A GPCR ^20^, is largely unchanged between the inactive and active-state SSTR2 conformations (Fig. 1f).

While the receptor activation mechanism appears the same between the two ligands, the extracellular receptor loops exhibit strikingly different behavior in the two structures. In the case of the 8-residue, SSTR2-selective octreotide, ECL2 folds down from its position in the apo state to form a cover over the ligand, making hydrogen bonds with the peptide and providing a cap involving W188 (Fig. 2a). ECL3 moves inward from the apo state and covers the sidechain of the N-terminal D-phenylalanine of octreotide with P286 (Extended Data Fig. 5a), providing hydrophobic packing. By contrast, the substantially larger SST14 peptide pushes ECL2 to be slightly more extended than in the apo state, with significant alterations in the pattern of residues interacting with the ligand (Fig. 2b). ECL3 is pulled inward towards the ligand in a similar overall fashion between SST14 and octreotide, although the P286 hydrophobic cap is in closer proximity with octreotide (Extended Data Fig. 5d). To probe the behavior of ECL2 we performed molecular dynamics (MD) simulations in triplicate at 1 μs timescale. The MD trajectories of apo-SSTR2 demonstrate that ECL2 remains in an upward state and does not spontaneously fold over the extracellular cavity (Fig. 2c), consistent with the cryo-EM structure. However, when simulations were performed starting from the apo state with octreotide added in the vicinity of the binding pocket, rapid repositioning of ECL2 was observed, demonstrating that this is a ligand-induced conformation of the receptor.

**Figure 2.**
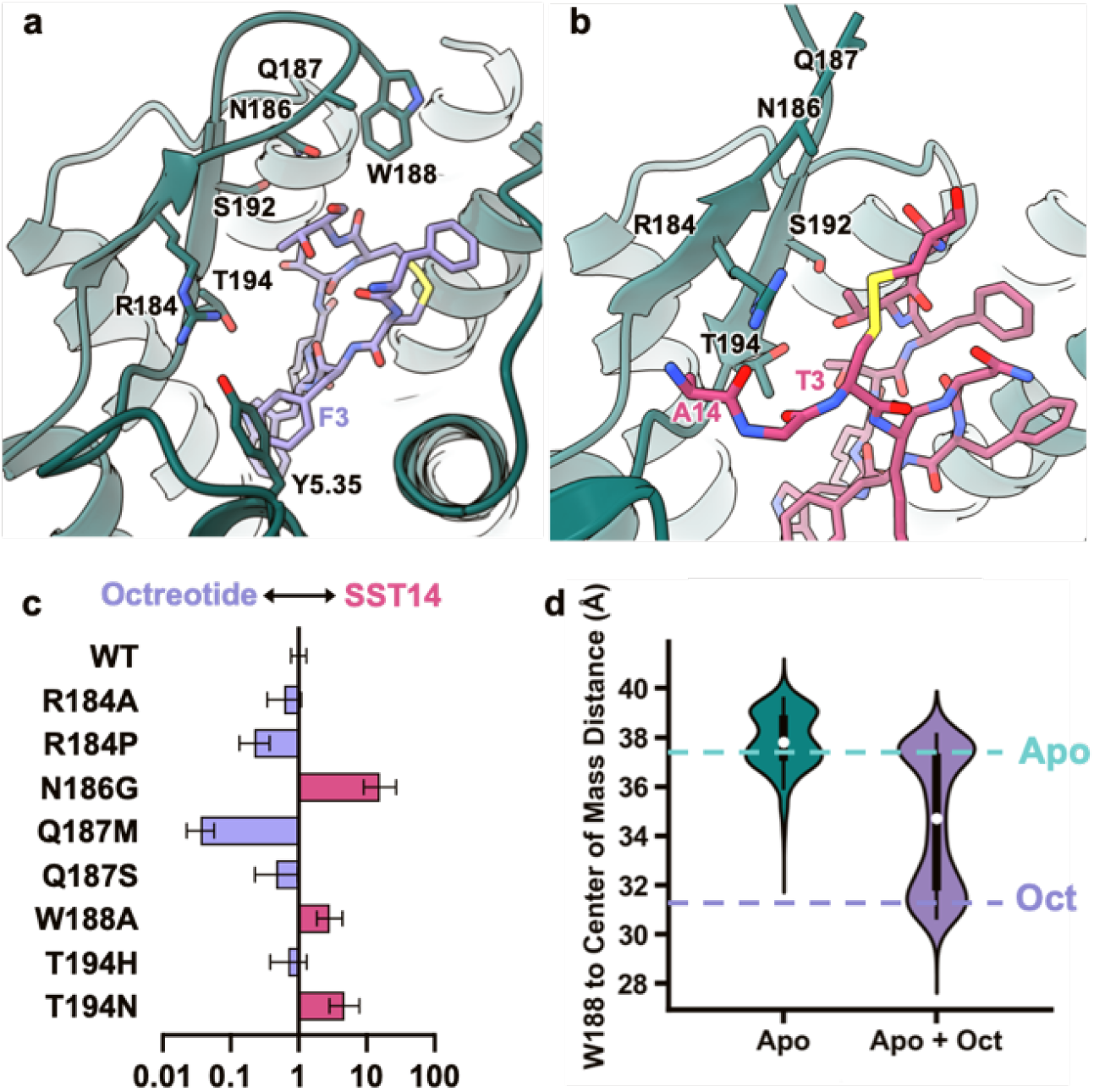
Differential SSTR2 ECL2 interactions with octreotide and SST14. **a,** ECL2 interactions of SST14 (magenta)-bound SSTR2 (teal). **b,** ECL2 interactions of octreotide (lavender)-bound SSTR2 (teal). **c,** Fold change in selectivity in favor of octreotide (lavender) or SST14 (magenta) as compared to selectivity of wild-type SSTR2. Error bars are 95% CI. **d,** Violin plots of the distance between W188 and the center of mass of the receptor during MD simulations of either apo inactive SSTR2 (teal) or apo SSTR2 with octreotide introduced (lavender).

To further explore the role of the ECL2-ligand interaction on agonist binding and consequent receptor activation, we mutated ECL2 residues to either alanine or analogous residues in other SSTR subtypes and evaluated their activity in Gi3 BRET-based assays (Fig. 2d, Extended Data Fig. 5 b, c). R184 is positioned to potentially form a hydrogen bond with the backbone of the SST14 N-terminus (Fig. 2b). In contrast, in the octreotidebound receptor, R184 forms an intra-receptor hydrogen bond with Y205^5.35^ (Fig. 2a), holding this residue in place to form an edge-face pi-pi interaction with F3 of octreotide. Mutation of R184 to either alanine or proline (analogous to SSTR1 and SSTR3, respectively) indeed substantially reduces SSTR2 activation by both compounds (Extended Data Fig. 5b, c). We also replaced T194 with either histidine or asparagine, present in SSTR3 and SSTR1, 4, & 5, respectively, which would disrupt both polar contacts made by T194 and provide a steric clash in the case of histidine. We found that both T194H and T194N impact the Gi3 signaling of both compounds (Extended Data Fig. 5b, c). N186 is observed near octreotide and positioned to make an indirect polar interaction with the ligand, while this residue is not resolved in the SST14 structure, presumably due to its variable positioning further up in the raised ECL2. Notably, mutation of N186 to glycine results in increased selectivity for SST14 over octreotide (Fig. 2d). On the other hand, Q187 at the tip of ECL2 does not appear to interact with either ligand directly, and the Q187S mutant retains identical Gi3 response and ligand selectivity compared to wild-type (Extended Data Fig. 5b, c). Curiously, the Q187M mutation enhances octreotide signaling (Fig. 2d), perhaps by generating additional hydrophobic packing with the neighboring W188. Finally, W188 occurs in an unresolved region in the SST14 structure but is resolved packed against octreotide’s C-terminal threonine-ol. Consistent with the notion that W188 participates only in octreotide binding, the W188G mutant altered selectivity to be unfavorable to octreotide (Fig. 2d).

### Subtype Selectivity at SSTRs is Multifaceted

The demonstration of ECL-driven ligand selectivity in SSTR2 raises questions about how the ECL2 and ECL3 of other SSTR subtypes contribute to ligand coordination and selectivity, as their ECL regions are highly divergent in length and sequence compared to SSTR2 (Fig. 3a). All SSTRs have comparable response to SST14 in BRET assays; however, octreotide activates SSTR2 most potently, with a reduced response at SSTR3 and SSTR5, and almost no activity at SSTR1 and SSTR4, which we recapitulated in our functional assays (Fig. 3b). To probe the role of individual SSTR ECLs on ligand recognition, we generated chimeric constructs of SSTR2 and used BRET-based assays to monitor Gi3 activation in response to SST14 and octreotide. In these chimeras we swapped the SSTR2 ECL2 or ECL3 for the corresponding sequence of other SSTR subtypes (Fig. 3a, c). For ECL2 we swapped the sequence either through the conserved cysteine (denoted “ECL2/X”) or in its entirety (denoted “ECL2+/X”, with X being the isoform number of origin). Surprisingly, nearly all ECL swaps impaired ligand recognition of SST14 and octreotide to some degree, with generally far greater impairment for SST14 than octreotide (Fig. 3d). Only the swap of the entirety of ECL2 from SSTR4 (ECL2+/4) selected against octreotide, and in this case, improved SST14-induced signaling compared to the shorter SSTR4 ECL2 swap. Further, exchanging ECL3 with the shorter ECL3/TM7 of SSTR1 and SSTR4 completely abolished G protein activation in response to either ligand.

**Figure 3.**
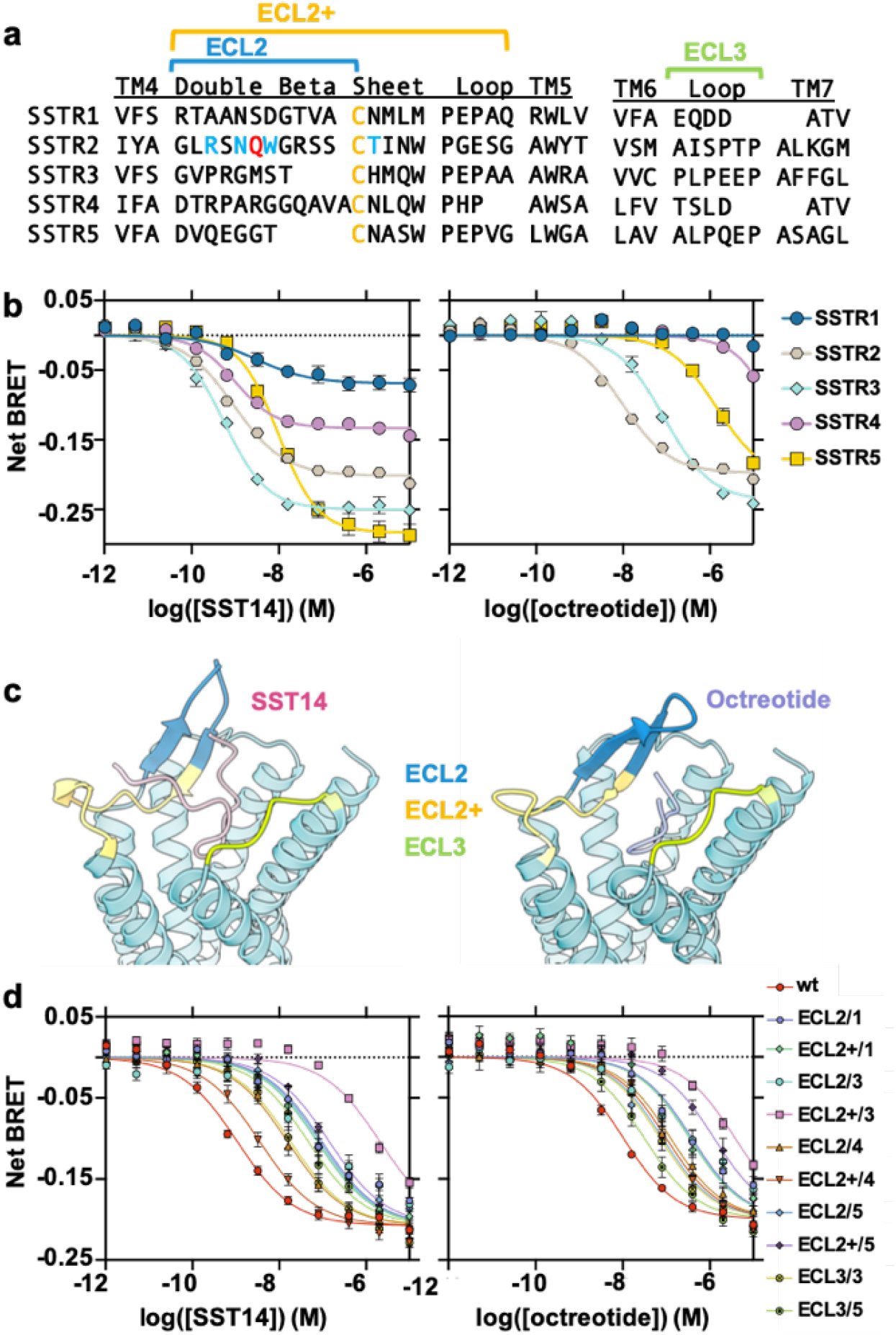
Role of ECL2 and ECL3 in SSTR subtype selectivity. **a,**Alignment of ECL2 and ECL3 of SSTRs, with swapped regions denoted with brackets.**b,**Dose-response curves of SSTR isoform-dependent activation of Gi3 BRET biosensor in response to SST14 (left) or octreotide (right). **c,** Structure of SST14 bound SSTR2 (left) and octreotide bound SSTR2 (right) with swapped regions colored. **d,** Dose-response curves of SSTR extracellular loop (ECL) swap-dependent activation of Gi3 BRET biosensor in response to SST14 (left) or octreotide (right). Error bars are *S.E.M.*.

The complexity of SST14 recognition can be further demonstrated through point mutations. H192 of SSTR3 is in a homologous position to T194 in SSTR2, which is located immediately next to the conserved ECL2 cysteine (C193 in SSTR2) that forms a disulfide bond with the conserved C^3.25^ (C115^3.25^ in SSTR2). Consistent with the observation that T194 in SSTR2 is critical for ligand recognition, we observe that H192T and H192A in SSTR3 are both deleterious to SST14-stimulated G protein activation (Extended Data Fig. 5f). Thus, despite the ECL2/C^3.25^ disulfide bond anchoring the position of this residue in space with respect to TM3 in all isoforms, mutation of the adjacent ECL2 T194 in SSTR2 or H192 in SSTR3 is functionally deleterious. Taken together, the mutagenesis and ECL swap results suggest that, whether through distinct peptide binding poses or via dynamic ECL interactions, endogenous SST14 forms divergent interactions with the extracellular portions of SSTRs.

Despite the complexity in ECL-driven receptor subtype selectivity, we were able to identify more straightforward sources of ligand discrimination in the TM bundle between SSTR1/4 and SSTR2/3/5. Prior work has suggested that mutating two SSTR1 TM bundle residues to the analogous residues of SSTR2, S305^7.35^F and Q291^6.55^N, can produce near-SSTR2 levels of octreotide binding^5^. Indeed, while SSTR2 but not SSTR1 is activated by octreotide, N276^6.55^Q in SSTR2 decreases octreotide signaling in a selective manner (Fig. 4c). This is consistent with our structural data, as the β-carbon of D-Trp in octreotide is only roughly 4 Å away from N276^6.55^ (Fig. 4d), clashing with a glutamine residue that would otherwise be accommodated by the L-Trp β-carbon of SST14. Furthermore, the effects of the S305^7.35^F mutation in SSTR1 (Fig. 4e) can be rationalized by considering F294^7.35^ in SSTR2, which lies below F6 in SST14 and the disulfide bond of octreotide (Fig. 4f), thus providing hydrophobic packing. Given the likelihood of significant conformational differences in peptide binding between subtypes, it seems plausible that the much larger phenylalanine of the S305^7.35^F mutation in SSTR1 would lock octreotide into a binding conformation akin to that of SSTR2. This model is consistent with prior studies that suggest the Q291^6.55^N alone does little to improve octreotide binding to SSTR1^5^. The SSTR2 structures also identify a novel source of SSTR1/4 versus SSTR2/3/5 selectivity. Although conserved between octreotide and SST14, T6/T10 (octreotide/SST14) is slightly twisted in octreotide compared to SST14, positioning this residue to interact with Q102^2.63^ (Fig. 4h). This glutamine, conserved in SSTR3 and 5 and replaced by a serine in SSTR1 and 4, appears important for ligand selectivity, as the Q102^2.63^S mutant in SSTR2 selectively reduces octreotide-induced G protein signaling (Fig. 4g).

**Figure 4.**
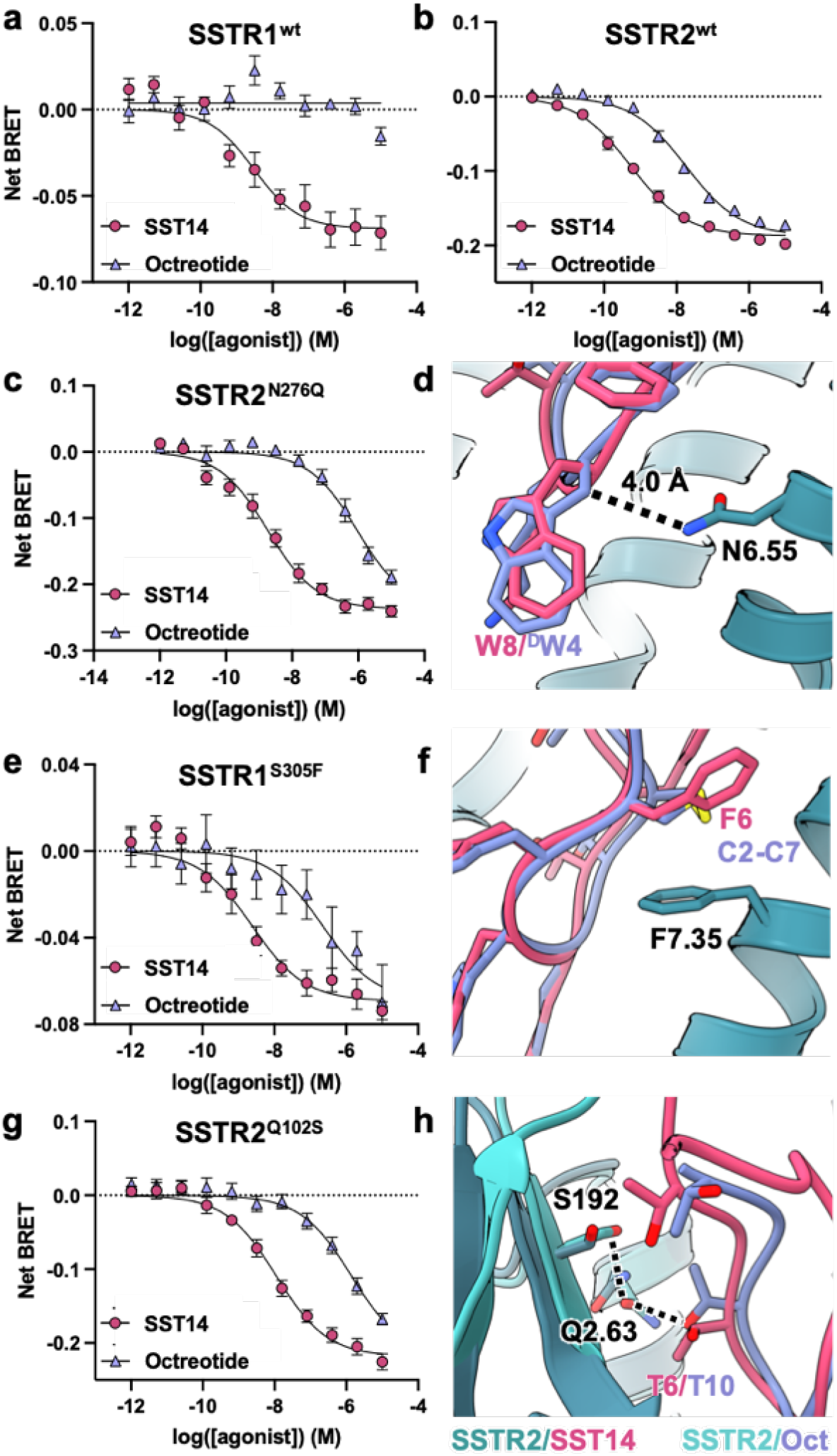
Subtype-selective point mutations in SSTR1 and SSTR2. Doseresponse curves for **a,** SSTR1- and **b,** SSTR2-dependent activation of Gi3 in response to SST14 and octreotide. **c,** Dose-response curves for SSTR2^N276Q^-dependent activation of Gi3, **d,** Overlay of SSTR2 (teal) bound to SST14 (magenta) and octreotide (lavender) highlighting the probable clash between octreotide D-Trp and a N6.55Q mutation as in SSTR1 and SSTR4. **e,** Dose-response curves for SSTR1^S305F^-dependent activation of Gi3, **f,** Overlay of SSTR2 (teal) bound to SST14 (magenta) and octreotide (lavender) highlighting the position of F7.35. **g,** Dose-response curves for SSTR2^Q102S^-dependent activation of Gi3, **h,** Overlay of SSTR2/SST14 structure (dark teal/magenta) with SSTR2/octreotide (light teal/lavender) highlighting differences in hydrogen bonding between T6/T10 of the ligands and ECL2/ Q102^2.63^. Error bars are *S.E.M.*.

## Discussion

In this study, we have presented structures of active-state SSTR2 in complex with either its native peptide agonist or a clinically used synthetic agonist. In combination with the structure of unliganded, inactive-state SSTR2 we report in an accompanying manuscript^12^, these structural snapshots provide a detailed view of the receptor activation mechanism and the interactions of the key tryptophan-lysine motif of SSTR peptide agonists. Our structural analysis and assays highlight a duality in the nature of subtype selectivity at SSTRs. The basis of selectivity in the TM bundles of SSTR1/4 versus SSTR2/3/5 is straightforward and largely attributable to three residues, yet complicated in the extracellular loops, where chimeric receptors have a non-linear effect on SST14-induced signaling. While it is clear from these studies that some aspects of subtype selectivity are structural, it is likely, given our discovery of the dynamic behavior of ECL2 and the observed non-linearity in ligand recognition of ECL chimeras, that some aspect of SSTR subtype selectivity is kinetically driven, as is the case for cannabinoid receptors^21^ and adrenergic receptors^22^. However, further structural, kinetic, and dynamics studies will be necessary to fully tease apart the intricacies of ligand recognition in this complex system. Despite this, the combination of high-resolution structures, precise characterization of activation mechanism, and a firm understanding of how residues in the transmembrane helices of SSTRs generate subtype selectivity provides a starting framework for the design of next-generation SSTR agonists.

## Supporting information

All Supplementary Figures

## Author Contributions

M.J.R. Cloned constructs, expressed and purified proteins, processed EM data, built models, and ran/analyzed molecular dynamics simulations. J.M. performed BRET assays, expressed and purified proteins. O.P. prepared cryo-EM samples and collected cryo-EM data. K.B. M.J.R. and G.S. wrote the manuscript with input from J.M. and O.P.. G.S. supervised the project.

## Data Availability

All data generated or analyzed in this study are included in this article and the Supplementary Information. The cryo-EM density maps and corresponding coordinates have been deposited in the Electron Microscopy Data Bank (EMDB) and the Protein Data Bank (PDB), respectively, under the following accession codes: XXXX YYYY

## Acknowledgements

Cryo-EM data were collected at the Stanford cryo-EM center (cEMc) with support from E. Montabana. This work was supported, in part, by the Mathers Foundation (G.S.), training grant T32GM089626 (J.G.M.) and used the Extreme Science and Engineering Discovery Environment (XSEDE)^23^ resource comet-gpu through sdsc-comet allocation TG-MCB190153 (G.S.), which is supported by National Science Foundation grant number ACI-1548562.

## Methods

### Construct generation

SSTR2 was obtained from Horizon Discovery in cDNA and cloned into a pFastBac vector containing an N-terminal haemagglutinin (HA) signal sequence followed by a FLAG epitope (DYKDDDDK), the 29 N-terminal residues (M1 to D29) of the β2 adrenergic receptor and TEV protease cleavage site; the C terminus contained a C3 protease cleavage site followed by enhanced green fluorescent protein (eGFP) and a hexahistine (His6) tag. The cloning was performed with Gibson cloning. The ICL3 and part of TM6 from Kappa opioid receptor was inserted as described in Robertson *et al*.^12^. Mutants and ECL swaps were generated with Q5 site directed mutagenesis.

### Expression and Purification of SSTR2-GFP

Baculovirus expressing SSTR2_KappaICL3_-GFP at P2 was used to infect Sf9 insect cells (Expression Systems) at a density of 3–4 million cells/ml and incubated at 28C with shaking. After 48 hours, cells were collected with centrifugation at 1500xg, washed with phosphate-buffered saline containing 2 μM agonist, and pellets were snap frozen in liquid nitrogen for purification. Pellets containing SSTR2_KappaICL3_-GFP were resuspended in hypotonic buffer containing 20 mM HEPES pH 7.5, 5 mM MgCl_2_, protease inhibitor cocktail, benzonase, 2 uM agonist (either octreotide or SST14), and 2 mg/ml iodoacetamide and gently stirred at 4C for an hour. Lysed membranes were harvested by centrifugation at 100,000xg. Membranes were resuspended in solubilization buffer containing 100 mM NaCl, 20 mM HEPES pH 7.5, protease inhibitor cocktail, 1 mM MgCl_2_, 10% glycerol, 2 uM agonist (either octreotide or SST14), and 2 mg/ml iodoacetamide and drip frozen in liquid nitrogen. On the day of purification, membranes were rapidly thawed and detergent was added dropwise to a final concentration of 1% LMNG, 0.2% CHS, 0.2% cholate and allowed to solubilize while gently stirring at 4C for 3 hours. Insoluble debris was removed with ultracentrifugation at 100,000xg, and solubilized receptor was supplemented with 20 mM imidazole and gravity loaded over Ni-NTA resin at 4C. The resin was washed with 10 column volume (CV) of buffer containing 100 mM NaCl, 20 mM HEPES pH 7.5, 2 uM agonist, 0.1% LMNG, 0.01% CHS, and 20 mM imidazole and protein was eluted in buffer consisting of 100 mM NaCl, 20 mM HEPES pH 7.5, 2 uM agonist, 0.01% LMNG, 0.001% CHS, 10% glycerol, and 250 mM imidazole. Eluted protein was supplemented with 5 mM CaCl_2_ and gravity loaded over M1 flag resin. The flag resin was washed with 10 CV of 100 mM NaCl, 20 mM HEPES pH 7.5, 2 uM agonist, 0.01% LMNG, 0.001% CHS, and 2 mM CaCl_2_. Protein was eluted in 100 mM NaCl, 20 mM HEPES pH 7.5, 2 uM agonist, 0.01% LMNG, 0.001% CHS, 10% glycerol, 0.1 mg/ml FLAG peptide and 2 mM EDTA, prior to concentration and injection onto size exclusion chromatography (SEC) with buffer consisting of 100 mM NaCl, 20 mM HEPES pH 7.5, 2 uM agonist, 0.01% LMNG, 0.001% CHS. Monomeric receptor was pooled, supplemented with glycerol, concentrated, and flash frozen in liquid nitrogen for later complexation.

### Expression and Purification of scFv16

The secreted single chain construct of Fab16 (scFv16) recognizes an epitope composed of the terminal part of the αN helix of Gαi1 as well as part of the Gβ1 subunit^13^ and was generated as previously described^15^. Purified scFv16 was concentrated to 13 mg/ml and flash frozen in buffer containing 20 mM HEPES pH 7.5, 100 mM NaCl, and 15% glycerol for later use.

### Purification of Gi protein heterotrimer

Pellets containing DN Gai3 were thawed and resuspended in lysis buffer containing 20 mM HEPES pH 75, 1 mM EDTA pH 8, 5% glycerol, 1 mM MgCl_2_, 5 uM β-mercaptoethanol, 100 uM GDP, protease inhibitor cocktail, and benzonase. Resuspended pellets were gently stirred for 20 minutes at 4C before pelleting membranes by ultracentrifugation at 100,000xg for 30 min. Membranes were resuspended with douncing in a glass tissue grinder in 100 mM NaCl, 20 mM HEPES pH 7.5, 1% sodium cholate, 5% glycerol, 1 mM MgCl_2_, 5 mM β-mercaptoethanol, 100 uM GDP, protease inhibitor cocktail, and benzonase. Solubilization further proceeded with 60 minutes of gentle stirring at 4C before insoluble debris was removed by ultracentrifugation at 100,000xg for 50 minutes. Solubilized G protein was supplemented with 30 mM imidazole and incubated with Ni-NTA beads for 1 hour. Beads were loaded onto a glass gravity column and washed with 10 CV with increasing concentrations of LMNG/CHS and decreasing concentrations of cholate until a final wash in 100 mM NaCl, 20 mM HEPES pH 7.5, 0.05% LMNG, 0.005% CHS, 5% glycerol, 1 mM MgCl_2_, 5 mM β-mercaptoethanol, 100 uM GDP, and 30 mM imidazole, before being eluted into the same buffer containing 250 mM imidazole. G protein heterotrimer was then supplemented with 1 mg HRV-3C protease/50 mg heterotrimer to remove the 6xHis tag and incubated overnight at 4C with dialysis against low imidazole buffer. The following day dialyzed protein was flowed through a Ni-NTA gravity column to remove HRV-3C and uncleaved heterotrimer, concentrated, and injected onto SEC with a buffer containing 100 mM NaCl, 20 mM HEPES pH 7.5, 0.01% LMNG, 0.001% CHS, 1 mM MgCl_2_, 100 uM TCEP, 20 uM GDP, and 5% glycerol. Fractions containing heterotrimer were pooled, concentrated, and flash frozen in liquid nitrogen for later use.

### Formation of SSTR2/Gi3 complex

Aliquots of SSTR2 _KappaICL3_-GFP and DN Gai3 were thawed and mixed in a buffer containing 100 mM NaCl, 20 mM HEPES pH 7.5, 0.01% LMNG, 0.001% CHS, 20 uM TCEP, 0.1mM MnCl2, lambda phosphatase, and 200 uM agonist peptide (either SST14 or octreotide), and incubated at room temperature for 1 hour. Complex was then incubated with 3C protease, apyrase, and scFV16 for 3 hours on ice. The mixture was then diluted 10-fold with buffer containing 100 mM NaCl, 20 mM HEPES pH 7.5, 0.01% LMNG, 0.001% CHS, 10 uM agonist peptide, and 5 mM CaCl_2_ and loaded onto M1 anti-DYKDDDDK immunoaffinity beads. Beads were washed with 10 CV of buffer containing 100 mM NaCl, 20 mM HEPES pH 7.5, 0.004% LMNG, 0.0004% CHS, 10 uM agonist peptide, and 1 mM CaCl_2_ to remove excess G protein and scFv before elution with 0.1 mg/ml FLAG peptide and 2 mM EDTA. Complex was concentrated and injected onto an enrich 650 SEC column equilibrated with buffer containing 100 mM NaCl, 20 mM HEPES pH 7.5, 0.001% LMNG, 0.00033% GDN, 0.0001% CHS, and 10 uM agonist peptide and fractions were spiked with 200 uM agonist peptide upon elution. Monomeric fractions were pooled, concentrated to 5-15 mg/ml, and used to freeze grids for cryogenic electron microscopy.

### Cryo-EM sample preparation and data collection

All samples were prepared on glow-discharged holey gold grids (Quantifoil ultrAufoil R1.2/1.3), blotted in an FEI Vitrobot Mark IV (Thermo Fisher Scientific) at 4C and 100% humidity, and plunge frozen into liquid ethane. Blotting conditions for each sample were as follows: 3.5 ul of SSTR2/SST14/Gi3/scFv16 complex at 15 mg/ml with an additional 0.025% beta OG, and 3.5 ul of SSTR2/octreotide/Gi3/scFv16 complex at 9 mg/ml.

Cryo-EM data were collected with a KRIOS electron microscope at an accelerating voltage of 300 kV using SerialEM with beam tilt compensation on a Gatan K3 direct electron detector. The resulting image stacks have a pixel size of 0.426 Å in super resolution mode. Each SSTR2/Gi3/Octreotide image stack is composed of 55 frames with an incident electron dose of 1.22 e-/Å2 per frame, for a total dose of 67 e-/Å2/s per micrograph. Each SSTR2/Gi3/SST14 image stack is composed of 50 frames with an incident electron dose of 1.04 e-/Å2 per frame, for a total dose of 52 e-/Å2/s per micrograph.

### Data Processing

All datasets were initially imported into Relion 3.1^24^ for motion correction with MotionCorr2^25^, CTF estimation with CTFFIND4^26^, and template-based particle picking. Extracted particles were then imported into CryoSPARC^27^ for 2D classification, 3D classification, and initial nonuniform refinement. Cleaned particle stacks were then transferred back to Relion 3.1 for Bayesian polishing, before being returned to CryoSPARC for final 2D cleaning, non-uniform, and local refinement. A pictorial flowchart of the data processing workflows can be found in Extended Data Figs. 2c, 3c.

### Model Building

Ligand placement was performed using a modification of the GemSpot pipeline^28^ to allow for placement of cyclic peptides. A custom library of ring conformations of the major ring was built for each peptide using PrimeMCS sampling^29^. This library was then used with GlideEM to produce a set of initial potential ligand poses. For each of these ligand poses, the full system, including protein, ligand and co-factors, was optimized using Phenix-OPLS^30^ to produce the final poses.

### Molecular Dynamics Simulations

The apo inactive state structure of SSTR2 determined in our accompanying manuscript was used to set up molecular dynamics simulations. Inactive receptor with octreotide was generated by aligning the active octreotide structure and translating the peptide to avoid steric clashes. Both systems were oriented with the OPM webserver^31^ in a lipid bilayer. The oriented systems were solvated in a box of POPC/CHS lipid bilayer, TIP3P water, and 150 mM NaCl with the CHARMM-GUI^32^. CHARMM36m^33^ input files with hydrogen mass repartitioning were taken from the CHARMM-GUI and used for MD simulations. The NAMD software package^34^ was used to execute simulations employing a Langevin thermostat and a Nosé-Hoover Langevin piston barostat at 1 atm with a period of 50 fs and decay of 25 fs; periodic boundary conditions were used with nonbonded interactions smoothed starting at 10 Å to 12 Å with long-range interactions treated with particle mesh Ewald (PME). A 2 fs timestep with SHAKE and SETTLE algorithms^35,36^ was used during equilibration and a 4 fs timestep during production. The system was minimized for 1,500 steps, heated from 0 to 303.15K in 20K increments simulating for 0.4 ns at each interval, and an additional 10 ns of equilibration was run at 10 ns; a 1 kcal/mol/Å2 harmonic restraint was applied to all non-hydrogen, non-water, and non-ion atoms for each of these steps. This was followed by 10 ns of equilibration with restraints applied to only non-hydrogen protein atoms, and then another 10 ns of equilibration with only CA atom restraints. 30 ns of unrestrained simulation was also considered to be equilibration; production simulations were performed for 1.0 μs. All simulations were run in triplicate with different initial velocity seeds for each condition. The distance between the center of mass of the receptor and the center of mass of W188 was measured in VMD^37^.

### JAWS calculations

JAWS simulation^17^ input files from the SSTR2/Gi3/SST14 complex structure were generated from a pdb file of all atoms within 25 A of the tip of the Gi3 C-terminal helix. The JAWS preparation scripts of the GemSpot pipeline^28^ were used to convert input files. A 15Å sphere around region of interest was solvated with theta waters to be sampled in the simulation and protein sidechains in this region were treated as flexible. The protein was simulated with the OPLS-AA/M force field^38^, and the TIP4P model^39^ was used for the water, with MCPRO^40^ used to for the JAWS Monte Carlo simulations. We ran 5 million steps for solvent equilibration, 10 million in hydration site identification, and 50 million in the production phase. Displayed waters are those where there was strong agreement between triplicate simulations in position and binding energy was estimated to be favorable (<0 kcal/mol).

### BRET-based assays

For BRET-based assays, SST14 (Cayman, 20809) was prepared in citrate buffer pH 4.8, octreotide (Cayman, 23757) in citrate buffer pH 4.8, neurotensin(8-13) (MedChem Express, HY-P0251) in water, and (-)-isoproterenol hydrochloride (Sigma, I6504) in water. All ligands were prepared at 10 mM, aliquoted, and stored at −80C for later use. Ligands used for cryoEM studies were prepared in water at a concentration of 10 mM. G protein BRET assays were performed as previously described (32367019) with the following modifications: HEK-293S cells grown in FreeStyle 293 suspension media (Thermo Fisher) were transfected at a density of 1 million cells/mL in 2 mL volume using 1200 ng total DNA at 1:1:1:1 ratio of receptor:G_α_:G_β_:G_γ_ and a DNA:PEI ratio of 1:5, and incubated in a 24 deep well plate at 220 rpm, 37°C for 48 hours. Cells were harvested by centrifugation, washed with Hank’s Balanced Salt Solution (HBSS) without Calcium/Magnesium (Gibco), and resuspended in assay buffer (HBSS with 20 mM HEPES pH 7.45) with 5 μg/mL freshly prepared coelenterazine 400a (GoldBio). Cells were then placed in white-walled, white-bottom 96 well plates (Costar) in a volume of 60 μl/well and 60,000 cells/well. Drug dilutions were prepared in drug buffer (assay buffer with 0.1% BSA, 6 mM CaCl_2_, 6 mM MgCl_2_), of which 30 μl were immediately added to plated cells. Ten minutes after the addition of ligand, plates were read using a SpectraMax iD5 plate reader using 585 nm and 525 nm emission filters with a one second integration time per well. The computed BRET ratios (GFP2/RLuc8 emission) were normalized to ligand-free control (Net BRET) prior to further analysis.

### Cell surface expression testing

Transfection of HEK-293S cells were performed with identical conditions as for BRET assays. Cells were washed with FACS buffer (1% BSA in PBS), stained with mouse anti-FLAG primary antibody (Sigma) in FACS buffer, washed three times, stained with Alexa Fluor 647 Goat Anti-Mouse IgG (Abcam), washed an additional three times with FACS buffer, and read on a NovoCyte Quanteon running NovoExpress v1.3.0 (Agilent) with >50,000 cells counted. Total expression levels (arbitrary units) were obtained via the product of percent positive by gating and median level of positive counts, and normalized to wild-type for at least 3 independent biological replicates.

### Statistical analysis

Normalized 11-point dose-response curves in technical duplicate were analyzed by simultaneous curve-fitting of at least 3 biological replicates (minimum 66 data points/curve) using a log(dose) vs. response model in Prism 9.1.0 for macOS (GraphPad Software) as previously described^10^. All 95% confidence intervals for EC_50_ and E_max_ were asymmetrically calculated, and S.E.M. of EC_50_ ratios were symmetrically calculated. All statistical comparisons between EC_50_ and EC_50_ ratios were performed with analysis of variance (ANOVA) (extra sum-of-squares F Test or one-way ANOVA) with correction for multiple hypothesis testing as previously described^10^.

## References

1. Günther, T. et al. International union of basic and clinical pharmacology. CV. somatostatin receptors: Structure, function, ligands, and new nomenclature. Pharmacol. Rev. (2018). doi:10.1124/pr.117.015388

2. Gu, Y. Z. & Schonbrunn, A. Coupling specificity between somatostatin receptor sst2A and G proteins: Isolation of the receptor-G protein complex with a receptor antibody. Mol. Endocrinol. (1997). doi:10.1210/mend.11.5.9926

3. Hofman, M. S., Eddie Lau, W. F. & Hicks, R. J. Somatostatin receptor imaging with68Ga DOTATATE PET/CT: Clinical utility, normal patterns, pearls, and pitfalls in interpretation1. Radiographics (2015). doi:10.1148/rg.352140164

4. Kaltsas, G., Androulakis, I. I., De Herder, W. W. & Grossman, A. B. Paraneoplastic syndromes secondary to neuroendocrine tumours. Endocrine-Related Cancer (2010). doi:10.1677/ERC-10-0024

5. Liapakis, G. et al. Identification of ligand binding determinants in the somatostatin receptor subtypes 1 and 2. J. Biol. Chem. (1996). doi:10.1074/jbc.271.34.20331

6. Bruns, C., Lewis, I., Briner, U., Meno-Tetang, G. & Weckbecker, G. SOM230: A novel somatostatin peptidomimetic with broad somatotropin release inhibiting factor (SRIF) receptor binding and a unique antisecretory profile. Eur. J. Endocrinol. (2002). doi: 10.1530/eje.0.1460707

7. Casarini, A. P. M. et al. Acromegaly: Correlation between expression of somatostatin receptor subtypes and response to octreotide-lar treatment. Pituitary (2009). doi: 10.1007/s11102-009-0175-1

8. Plöckinger, U., Dienemann, D. & Quabbe, H.-J. Gastrointestinal Side-Effects of Octreotide during Long Term Treatment of Acromegaly*. J. Clin. Endocrinol. Metab. (1990). doi:10.1210/jcem-71-6-1658

9. Parry, J. J., Chen, R., Andrews, R., Lears, K. A. & Rogers, B. E. Identification of critical residues involved in ligand binding and G protein signaling in human somatostatin receptor subtype 2. Endocrinology (2012). doi:10.1210/en.2011-1662

10. Olsen, R. H. J. et al. TRUPATH, an open-source biosensor platform for interrogating the GPCR transducerome. Nat. Chem. Biol. (2020). doi: 10.1038/s41589-020-0535-8

11. Che, T. et al. Nanobody-enabled monitoring of kappa opioid receptor states. Nat. Commun. (2020). doi:10.1038/s41467-020-14889-7

12. Robertson, M. J. et al. Structure Determination of Inactive-State GPCRs with a Universal Nanobody. (2021).

13. Maeda, S. et al. Development of an antibody fragment that stabilizes GPCR/G-protein complexes. Nat. Commun. (2018). doi:10.1038/s41467-018-06002-w

14. Maeda, S., Qu, Q., Robertson, M. J., Skiniotis, G. & Kobilka, B. K. Structures of the M1 and M2 muscarinic acetylcholine receptor/G-protein complexes. Science (80-.). 364, 552–557 (2019).

15. Koehl, A. et al. Structure of the μ-opioid receptor-Gi protein complex. Nature 558, 547–552 (2018).

16. Ballesteros, J. A. & Weinstein, H. Integrated methods for the construction of three-dimensional models and computational probing of structure-function relations in G protein-coupled receptors. Methods Neurosci. (1995). doi: 10.1016/S1043-9471(05)80049-7

17. Michel, J., Tirado-Rives, J. & Jorgensen, W. L. Prediction of the water content in protein binding sites. J. Phys. Chem. B 113, 13337–13346 (2009).

18. Wacker, D. et al. Structural features for functional selectivity at serotonin receptors. Science (80-.). (2013). doi:10.1126/science.1232808

19. Deupi, X., Standfuss, J. & Schertler, G. Conserved activation pathways in G-protein-coupled receptors. Biochemical Society Transactions (2012). doi:10.1042/BST20120001

20. McAllister, S. D. et al. Structural Mimicry in Class A G Protein-coupled Receptor Rotamer Toggle Switches. J. Biol. Chem. (2004). doi:10.1074/jbc.m406648200

21. Xing, C. et al. Cryo-EM Structure of the Human Cannabinoid Receptor CB2-Gi Signaling Complex. Cell (2020). doi:10.1016/j.cell.2020.01.007

22. Xu, X. et al. Binding pathway determines norepinephrine selectivity for the human β1AR over β2AR. Cell Res. (2021). doi:10.1038/s41422-020-00424-2

23. Towns, J. et al. XSEDE: Accelerating scientific discovery. Comput. Sci. Eng. (2014). doi:10.1109/MCSE.2014.80

24. Zivanov, J. et al. RELION-3: New tools for automated high-resolution cryo-EM structure determination. bioRxiv (2018). doi:10.1101/421123

25. Zheng, S. Q. et al. MotionCor2: Anisotropic correction of beam-induced motion for improved cryo-electron microscopy. Nature Methods 14, 331–332 (2017).

26. Rohou, A. & Grigorieff, N. CTFFIND4: Fast and accurate defocus estimation from electron micrographs. J. Struct. Biol. 192, 216–221 (2015).

27. Punjani, A., Rubinstein, J. L., Fleet, D. J. & Brubaker, M. A. CryoSPARC: Algorithms for rapid unsupervised cryo-EM structure determination. Nat. Methods (2017). doi:10.1038/nmeth.4169

28. Robertson, M. J., van Zundert, G. C. P., Borrelli, K. & Skiniotis, G. GemSpot: A Pipeline for Robust Modeling of Ligands into Cryo-EM Maps. Structure (2020). doi:10.1016/j.str.2020.04.018

29. Sindhikara, D. et al. Improving Accuracy, Diversity, and Speed with Prime Macrocycle Conformational Sampling. J. Chem. Inf. Model. (2017). doi: 10.1021/acs.jcim.7b00052

30. van Zundert, G. C. P., Moriarty, N. W., Sobolev, O. V., Adams, P. D. & Borrelli, K. W. Macromolecular refinement of X-ray and cryoelectron microscopy structures with Phenix/OPLS3e for improved structure and ligand quality. Structure (2021). doi:10.1016/j.str.2021.03.011

31. Lomize, M. A., Pogozheva, I. D., Joo, H., Mosberg, H. I. & Lomize, A. L. OPM database and PPM web server: Resources for positioning of proteins in membranes. Nucleic Acids Res. 40, (2012).

32. Lee, J. et al. CHARMM-GUI Input Generator for NAMD, GROMACS, AMBER, OpenMM, and CHARMM/OpenMM Simulations Using the CHARMM36 Additive Force Field. J. Chem. Theory Comput. 12, 405–413 (2016).

33. Huang, J. et al. CHARMM36m: An improved force field for folded and intrinsically disordered proteins. Nat. Methods 14, 71–73 (2016).

34. Phillips, J. C. et al. Scalable molecular dynamics with NAMD. Journal of Computational Chemistry 26, 1781–1802 (2005).

35. Miyamoto, S. & Kollman, P. A. Settle: An analytical version of the SHAKE and RATTLE algorithm for rigid water models. J. Comput. Chem. (1992). doi:10.1002/jcc.540130805

36. Ryckaert, J. P., Ciccotti, G. & Berendsen, H. J. C. Numerical integration of the cartesian equations of motion of a system with constraints: molecular dynamics of n-alkanes. J. Comput. Phys. (1977). doi:10.1016/0021-9991(77)90098-5

37. Humphrey, W., Dalke, A. & Schulten, K. VMD: Visual molecular dynamics. J. Mol. Graph. 14, 33–38 (1996).

38. Robertson, M. J., Tirado-Rives, J. & Jorgensen, W. L. Improved Peptide and Protein Torsional Energetics with the OPLS-AA Force Field. J. Chem. Theory Comput. 11, 3499–3509 (2015).

39. Jorgensen, W. L., Chandrasekhar, J., Madura, J. D., Impey, R. W. & Klein, M. L. Comparison of simple potential functions for simulating liquid water. J. Chem. Phys. 79, 926–935 (1983).

40. Jorgensen, W. L. & Tirado-Rives, J. Molecular modeling of organic and biomolecular systems using BOSS and MCPRO. J. Comput. Chem. 26, 1689–1700 (2005).

