## Supplementary material for "Plasticity in Ligand Recognition at Somatostatin Receptors": All Supplementary Figures

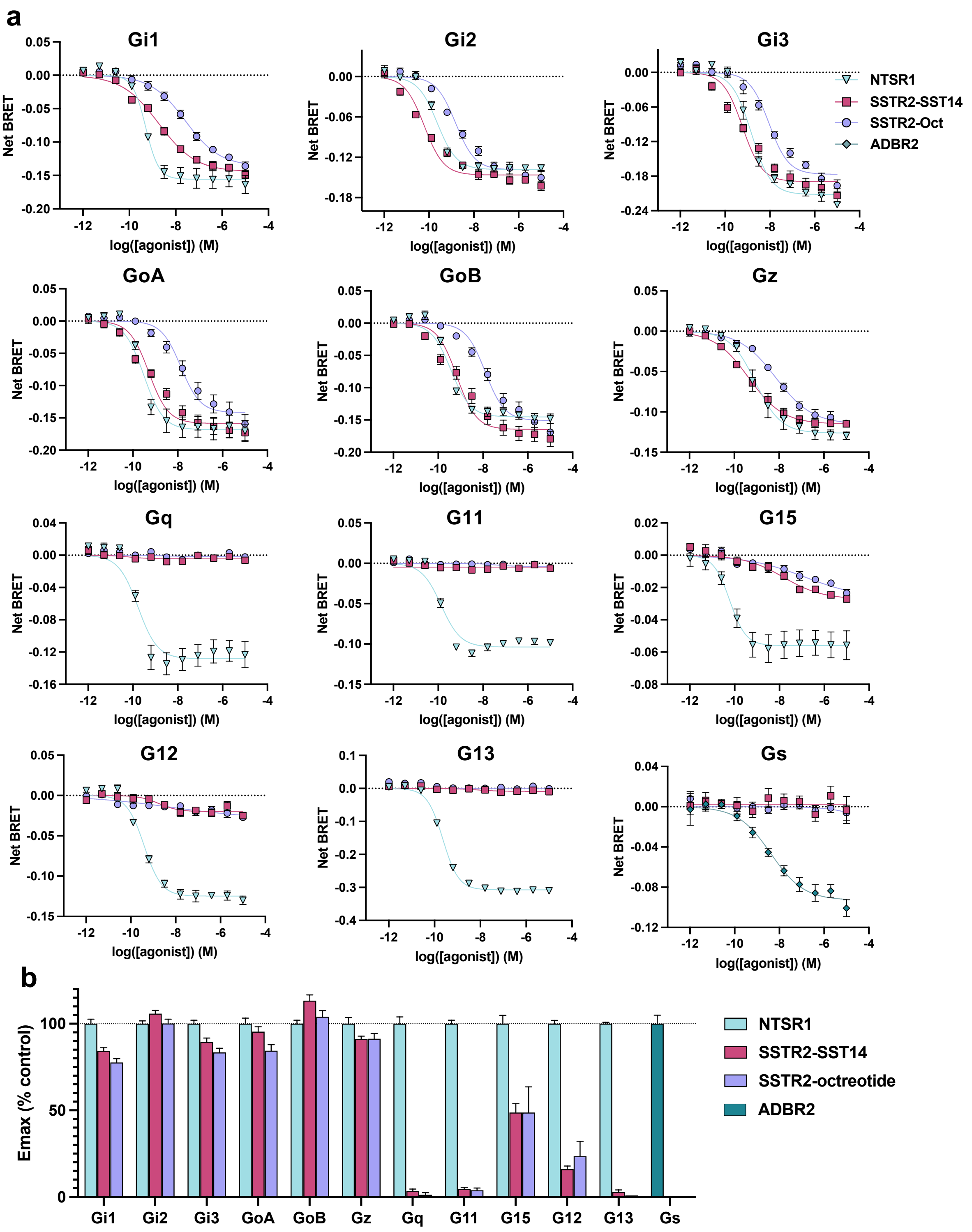


**Extended Data Fig. 1 | G protein specificity of SSTR2. a**, Dose-response curves of G protein-specific activation pathways of SSTR2 by SST14 (magenta squares), or octreotide (lavender circles), as compared to activation of neurotensin 1 receptor (NTSR1) by neurotensin (blue triangles) or $\beta$2 adrenergic receptor by isoproterenol (teal diamonds) from simultaneous curve fitting of 3 independent biological replicates with Hill Slope constrained to 1. Error bars are *S.E.M.*. **b,** Bar plot of $E_{max}$ from the above dose-response curves. Error bars are *S.E.M.*.

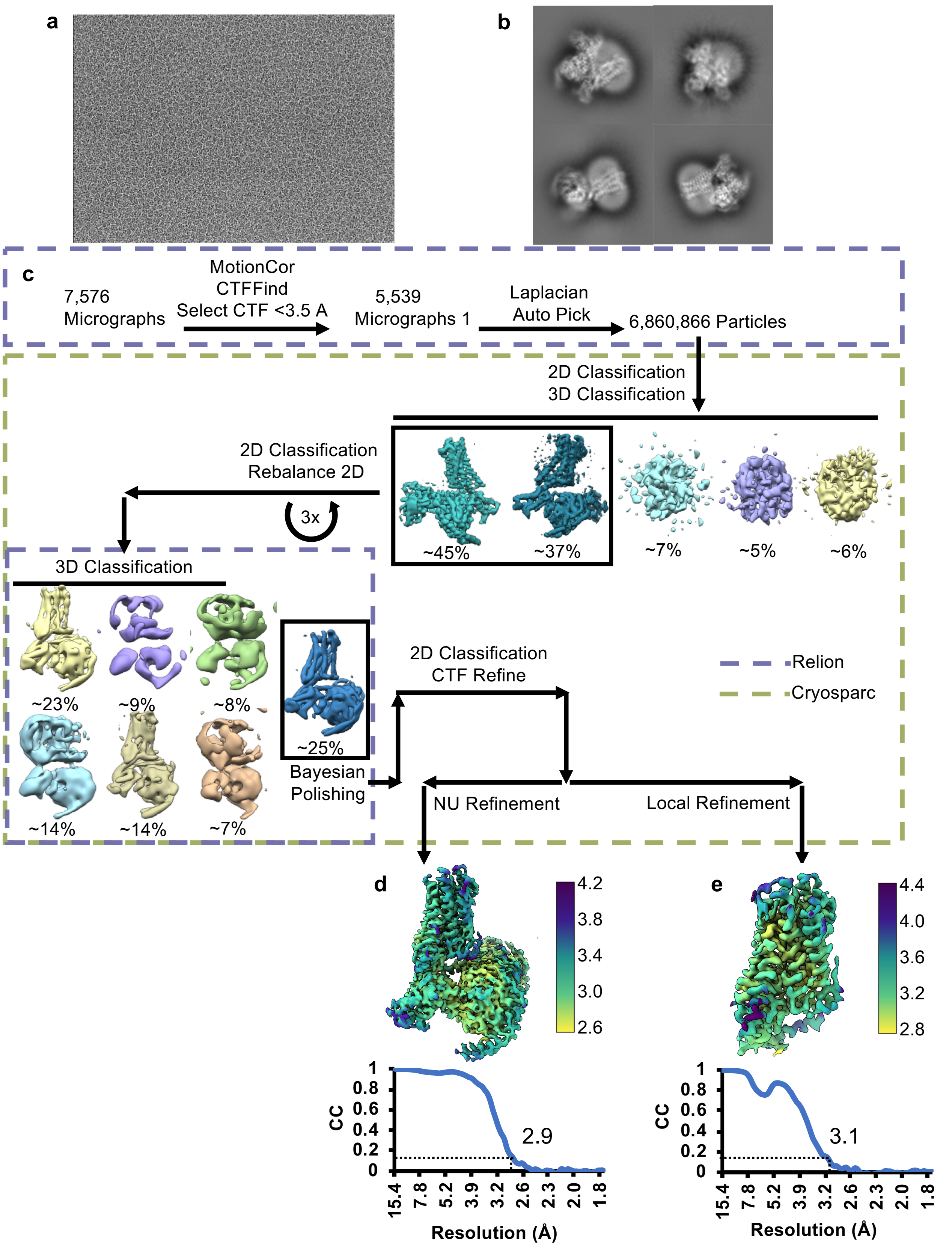


**Extended Data Fig. 2 | SSTR2/Gi3/scFv16/octreotide complex cryo-EM data collection and processing. a,** Representative micrograph of SSTR2/Gi3/scFv16/octreotide complex. **b,** Example final 2D classes of SSTR2/Gi3/scFv16/octreotide complex. **c,** Cryo-EM data processing workflow. **d,** Local resolution of SSTR2/Gi3/scFv16/octreotide global refinement with FSC curve below. **e,** Local resolution of SSTR2/Gi3/scFv16/octreotide local refinement on SSTR2 with FSC curve below.

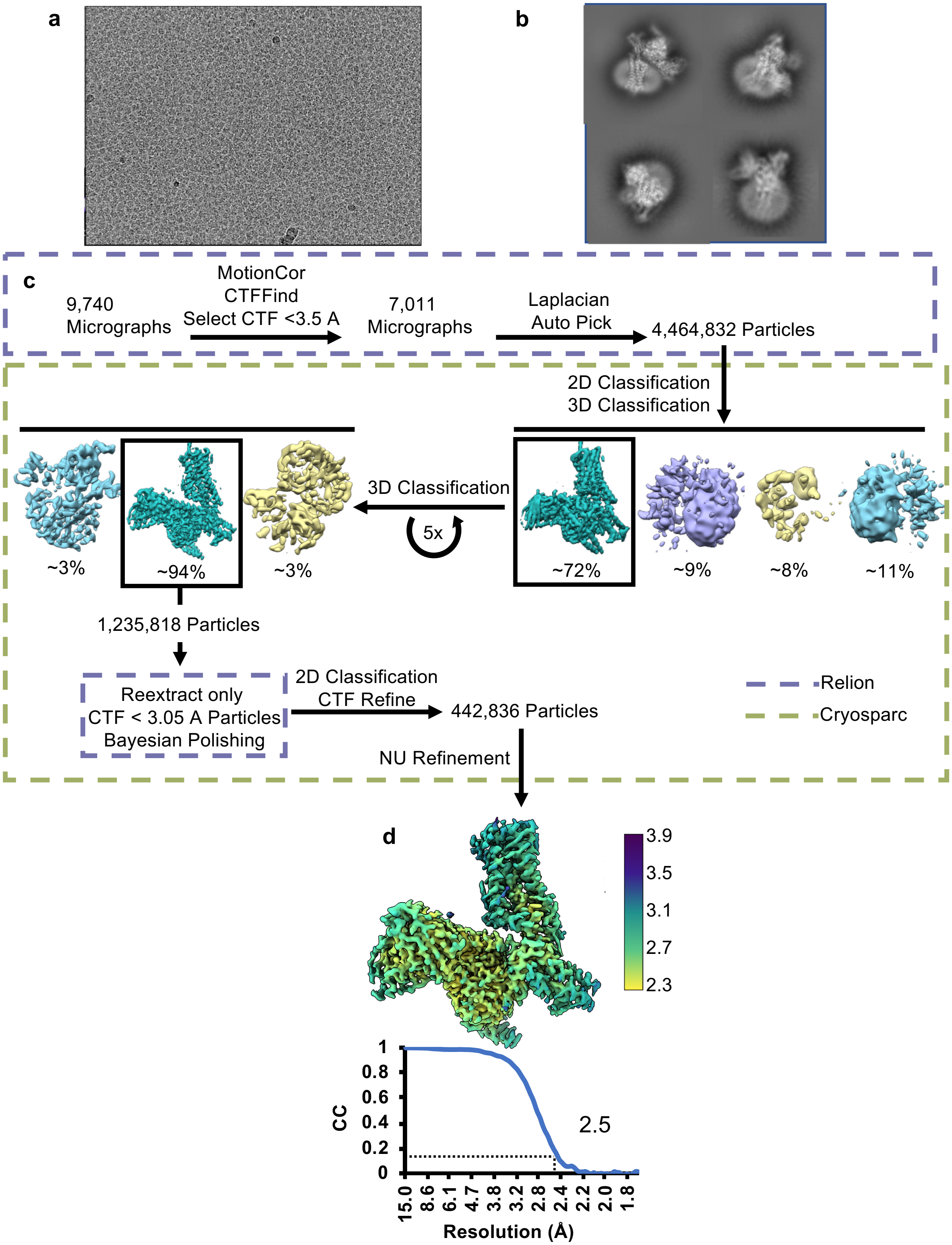


**Extended Data Fig. 3 | SSTR2/Gi3/scFv16/SST14 complex cryo-EM data collection and processing. a,** Representative micrograph of SSTR2/Gi3/scFv16/SST14 complex. **b,** Example final 2D classes of SSTR2/Gi3/scFv16/SST14 complex. **c,** Cryo-EM data processing workflow. **d,** Local resolution of SSTR2/Gi3/scFv16/SST14 global refinement with FSC curve below.

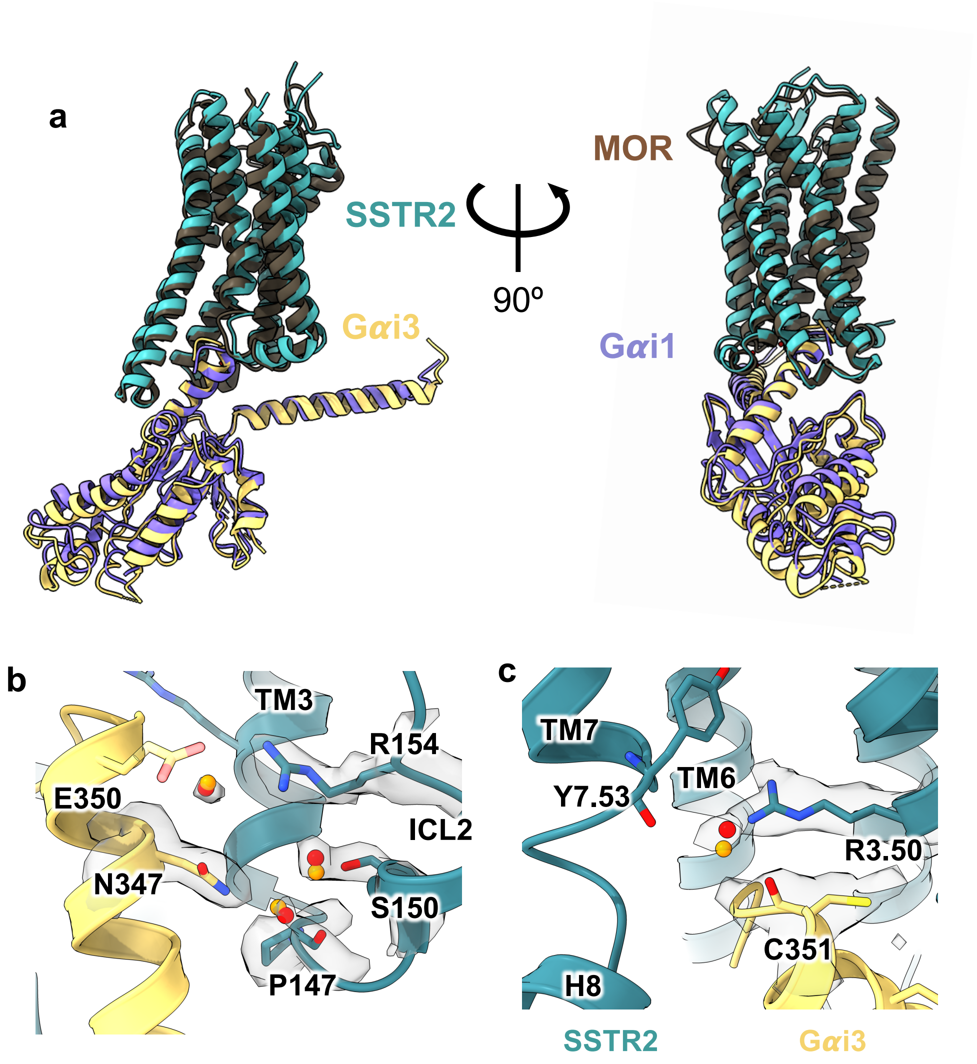


**Extended Data Fig. 4 | Comparison of SSTR2 Gi3 interface. a,** Alignment of SSTR2/Gi3 and MOR Gi1 from two different angles. **b,** Hydration at the SSTR2 ICL2/Gi3 interface; orange spheres are predicted water positions from JAWS simulations. **c,** Hydration at the SSTR2 DRY motif/Gi3 interface; orange spheres are predicted water positions from JAWS simulations.

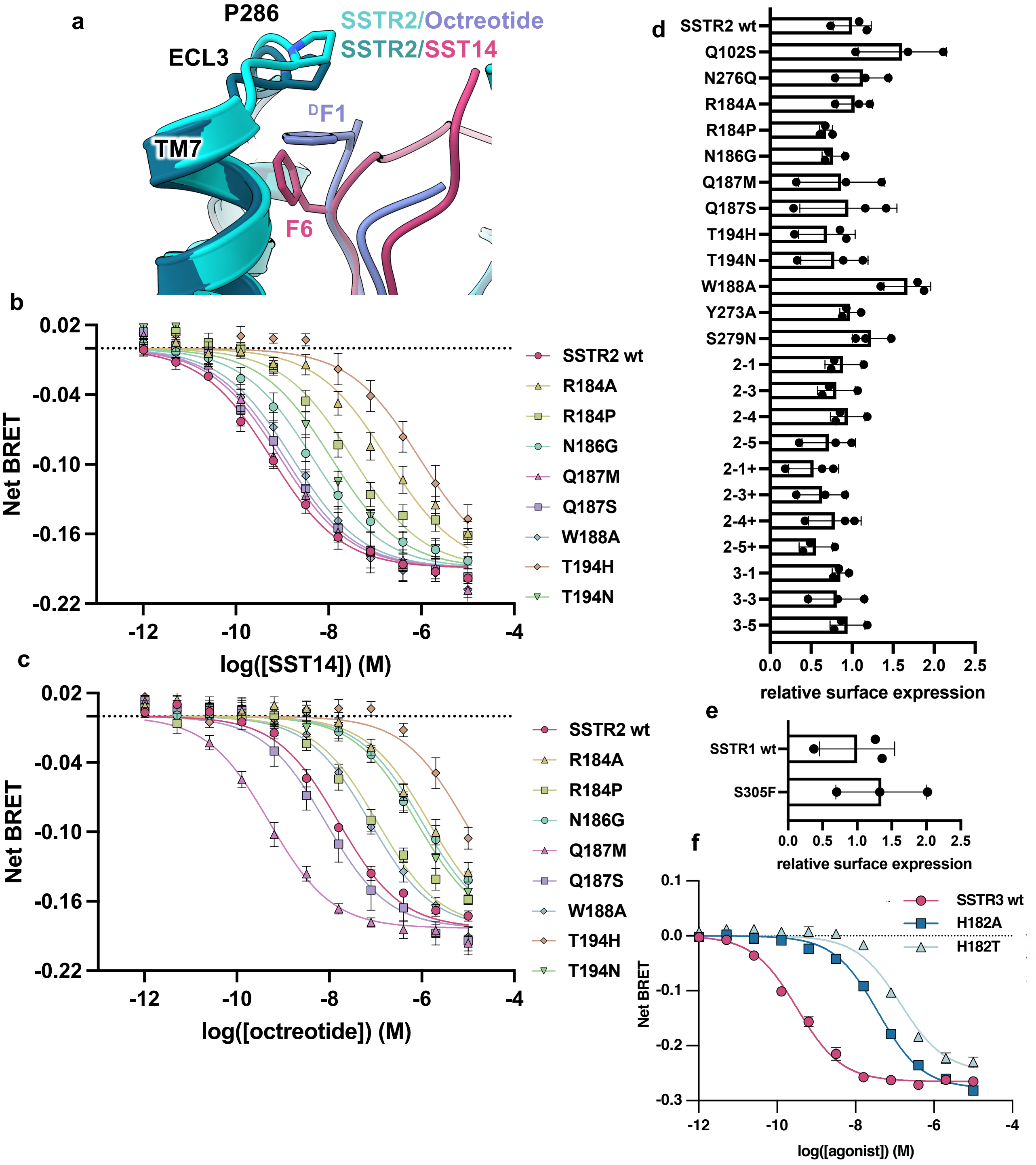


**Extended Data Fig. 5 | Comparison of ECL2 and ECL3 interactions and mutagenesis with octreotide and SST14. a,** Overlay of the structures of SSTR2 bound to octreotide (lavender) and SST14 (magenta) around ECL3. **b,** Dose-response curves of SSTR2-dependent Gi3 BRET biosensor activation by SST14. **c,** Dose-response curves of SSTR2-dependent Gi3 BRET biosensor activation by octreotide. Error bars are *S.E.M.*. **d,** Cell surface expression analysis of point mutants and ECL swaps of SSTR2 and **e,** SSTR1. **f,** Dose-response curves of SSTR3-dependent Gi3 BRET biosensor activation by SST14. Statistical test performed using one-way ANOVA with Bonferroni correction. Error bars are *S.D.*.

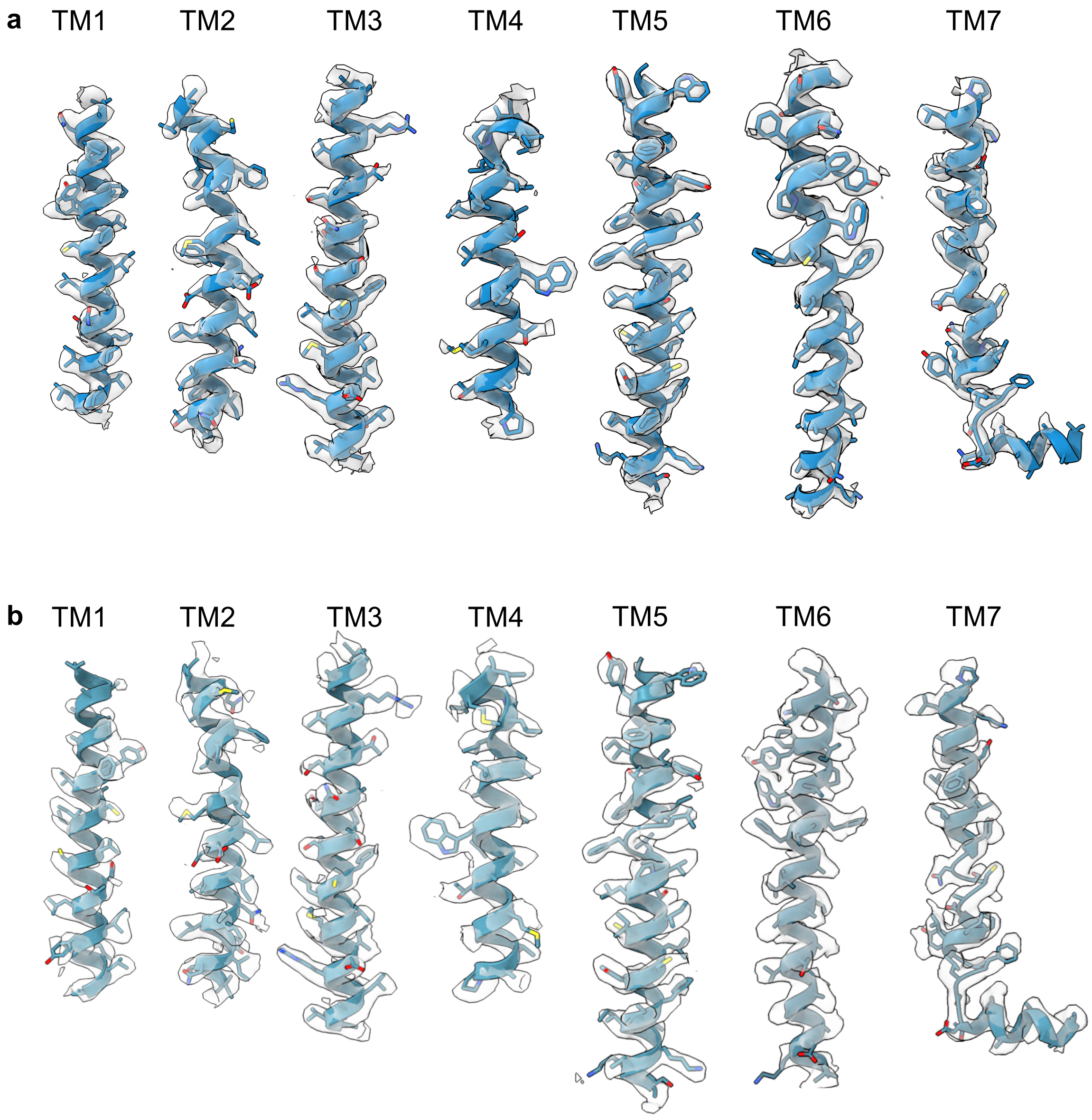


**Extended Data Fig. 6 | Map-Model Agreements. a,** Map-model comparison for SSTR2/SST14. **b,** Map-model comparison for SSTR2/Octreotide.

**Extended Data Table 1 | Cryo-EM data collection, refinement and validation statistics**

| | SSTR2/Octreotide/ Gi3/scFv16 | SSTR2/SST14/ Gi3/scFv16 |
|---|---|---|
| **Data collection and processing** | | |
| Magnification (x) | 57,050 | 57,050 |
| Voltage (kV) | 300 | 300 |
| Electron exposure (e–/Å2) | 67.00 | 52.00 |
| Defocus range (μm) | -0.8 to -1.8 | -0.8 to -1.8 |
| Pixel size (Å) | 0.8521 | 0.8521 |
| Symmetry imposed | C1 | C1 |
| Initial particle images (no.) | 6,860,866 | 4,464,832 |
| Final particle images (no.) | 281,479 | 442,863 |
| Map resolution (Å) | 2.9 | 2.5 |
| FSC threshold | 0.143 | 0.143 |
| **Refinement** | | |
| Model Resolution | 2.9 | 2.5 |
| FSC Threshold | 0.143 | 0.143 |
| *Model Composition* | | |
| Non-hydrogen Atoms | 8215 | 8428 |
| Protein Atoms | 8215 | 8423 |
| Waters & Ions | 0 | 5 |
| *B factor (Å$^2$)* | | |
| Protein Atoms | 46.84 | 52.86 |
| Waters & Ions | | 49.20 |
| *R.M.S Deviations* | | |
| Bonds (Å) | 0.005 | 0.005 |
| Angles (°) | 0.907 | 0.997 |
| *Validation* | | |
| MolProbity score | 1.58 | 1.33 |
| Clashscore | 5.26 | 3.49 |
| Poor rotamers (%) | 0.13 | 0.73 |
| *Ramachandran Plot* | | |
| Favored (%) | 95.69 | 96.89 |
| Allowed (%) | 4.31 | 3.02 |
| Outliers (%) | 0.00 | 0.09 |
